# Beyond type 1 regulatory T cells: co-expression of LAG3 and CD49b in IL-10-producing T cell lineages

**DOI:** 10.1101/359547

**Authors:** Weishan Huang, Sabrina Solouki, Chavez Carter, Song-Guo Zheng, Avery August

## Abstract

Type 1 regulatory CD4^+^ T (Tr1) cells express high levels of the immunosuppressive cytokine IL-10 but not the master transcription factor Foxp3, and can suppress inflammation and promote immune tolerance. In order to identify and obtain viable Tr1 cells for research and clinical applications, co-expression of CD49b and LAG3 has been proposed as a unique surface signature for both human and mouse Tr1 cells. However, recent studies have revealed that this pattern of co-expression is dependent on the stimulating conditions and the differentiation stage of the CD4^+^ T cells. Here, using an IL-10^GFP^/Foxp3^RFP^ dual reporter transgenic murine model, we demonstrate that co-expression of CD49b and LAG3 is not restricted to the Foxp3^−^ Tr1 cells, but is also observed in Foxp3^+^ T regulatory (Treg) cells and CD8^+^ T cells that produce IL-10. Our data indicate that IL-10-producing Tr1 cells, Treg cells and CD8^+^ T cells are all capable of co-expressing LAG3 and CD49b *in vitro* following differentiation under IL-10-inducing conditions, and *in vivo* following pathogenic insult or infection in the pulmonary mucosa. Our findings urge caution in the use of LAG3/CD49b co-expression to identify Tr1 cells, since it may mark IL-10-producing T cell lineages more broadly, including the Foxp3^−^ Tr1 cells, Foxp3^+^ Treg cells and CD8^+^ T cells.

## INTRODUCTION

The mammalian immune system has evolved both effector and regulatory immune axes to protect the host from invading pathogens, along with a control mechanism to tune the level of immune reactivity against self- and non-self-agents to prevent host tissue damage. Interleukin-10 (IL-10) is a regulatory cytokine with demonstrated anti-inflammatory function and plays an essential role in preventing allergic inflammation [1], autoimmunity [2] and pathogen-induced immunopathology [3; 4], but can also promote the establishment and maintenance of chronic infection [5; 6]. IL-10 has been reported as a product of activation of multiple immune cell lineages. Innate immune cells including dendritic cells (DCs) [7], macrophages [8], neutrophils [9] and innate lymphoid cells (ILCs) [10] have been reported to express IL-10 *in vivo* and *in vitro*. IL-10 is also expressed by many subsets of the adaptive immune cells, including B cells [11] and T cells including the Foxp3^−^ CD4^+^ [12], Foxp3^+^ Treg [13] and CD8^+^ T cell subsets [14]. Regulatory T cells are defined by their immunosuppressive function, and the three aforementioned subsets of IL-10-producing T cells have been reported as phenotypically distinct regulatory T cell subsets, playing important roles in promoting immune tolerance and/or suppressing inflammation in both mouse and human [15; 16; 17; 18; 19; 20].

Among the IL-10-producing T cells, the Foxp3^−^ CD4^+^ T cell subset, also known as type 1 regulatory T cells (Tr1 cells), are inducible in the periphery and have a pivotal role in limiting inflammation [15; 21; 22; 23]. Tr1 cells have been shown to prevent allergic asthma [24] and atopic dermatitis [25] in murine models. In both mouse models and humans, induction of tolerance *via* specific antigen immunotherapy (SIT) is accompanied by induction of Tr1 cells [26; 27]. Therefore, Tr1 cells have strong promise as a potential therapeutic approach for inflammatory diseases. Tr1 cells can be differentiated from naïve CD4^+^ T cells upon TCR engagement in the presence of IL-27 *in vitro* [28], and in order to identify and obtain viable Tr1 cells for clinical application, co-expression of LAG3 and CD49b has been recently proposed to be cell surface signature of the Foxp3^−^ IL-10^high^ Tr1 cells [15]. LAG3 is a structural homolog of CD4 molecule and can bind to MHC class II with high affinity [29; 30]. LAG3 is highly expressed by IL-10^+^ CD4^+^ T cells [31], as well as by activated effector T cells [32] and Foxp3^+^ Treg cells [33]. CD49b is the α2 integrin subunit, highly expressed by NK cells [34]. CD49b is up-regulated in T cells that may produce IL-10 and/or pro-inflammatory cytokines [35; 36; 37]. In addition to Foxp3^−^ Tr1 cells, IL-10 can be highly up-regulated in activated Foxp3^+^ Treg and CD8^+^ T cells under inflammatory conditions and/or upon TCR activation. Given the importance of being able to identify Foxp3^−^ Tr1 cells, including under clinical conditions, and to gain a better understanding of the specificity of co-expression of LAG3 and CD49b as a cell surface signature for IL-10-producing cells, we sought to determine whether co-expression of LAG3 and CD49b can mark broader range of T cell subsets that are actively producing high levels of IL-10.

Using a murine model carrying an IL-10^GFP^/Foxp3^RFP^ dual reporter system, we find that co-expression of LAG3 and CD49b is a generic feature of the IL-10-producing Foxp3^−^ CD4^+^, Foxp3^+^ CD4^+^ and CD8^+^ T cell subsets. The capacity of co-expression of LAG3 and CD49b expression in marking IL-10^high^ T cell subsets is dependent on the disease conditions and anatomical locations of the cells. Furthermore, co-expression of LAG3 and CD49b expression is also a shared feature of human IL-10-producing FOXP3^−^ CD4^+^, FOXP3^+^ CD4^+^ and CD8^+^ T cell subsets. Our data reveal that co-expression of LAG3 and CD49b is a generic signature of IL-10-producing T cells, which is broader than previously appreciated.

## MATERIALS AND METHODS

### Mice and human blood samples

All mice were on the C57BL/6 background. *Rag1^−/−^* (B6.129S7-*Rag1^tm1Mom^*/J), IL-10^GFP^ (B6(Cg)-*Il10^tm1.1Karp^*/J) [38], and Foxp3^RFP^ (C57BL/6-*Foxp3^tm1Flv^*/J) [39] reporter mice were from the Jackson Laboratory (Bar Harbor, ME). Single reporter strains were crossed to generate an IL-10^GFP^/Foxp3^RFP^ dual reporter strain as we recently reported [40]. Human peripheral blood samples were procured from the New York Blood Center collected from healthy cohorts. All experiments were approved by the Office of Research Protection’s Institutional Animal Care and Use Committee and Institutional Review Board at Cornell University.

### Antibodies and other reagents

All fluorescent antibodies are listed in “fluorochrome-target (clone; annotation if desirable)” format below.

#### Mouse antibodies

Purified anti-CD16/32 (93; Fc block), CD3ε (145-2C11), CD28 (37.51), IFN-γ(XMG1.2), and IL-12 (C17.8) antibodies were from BioLegend (San Diego, CA); Pacific Blue-CD90 (53-2.1), FITC-TCRβ (H57-597), APC-LAG3 (C9B7W), PE-Cy7-CD49b (HMγ2), and PE-Cy7-CD62L (MEL-14) were from BioLegend; eFluor 450-CD4 (GK1.5); Alexa Fluor 700-CD4 (GK1.5) were from eBioscience; BD Horizon V500-CD44 (IM7), PE-CD44 (IM7), and APC-Cy7-TCRβ (H57-597) were from BD Biosciences; PerCP-Cy5.5-CD8α (2.43) was from Tonbo Biosciences.

#### Human antibodies

Purified anti-CD3ε (OTK3) and CD28 (28.2), eFluor 450-CD8α (RPA-T8) FITC-CD4 (OKT4), and APC-FOXP3 (236A/E7) were from eBioscience; PE-IL-10 (JES3-19F1), Alexa Fluor 647-LAG3 (11C3C65), and PerCP-Cy5.5-LAG3 (11C3C65) were from BioLegend; FITC-CD49b (AK-7) and Alexa Fluor 700-CD4 (RPA-T4) were from BD Biosciences.

#### Other reagents

Human TruStain FcX (Fc receptor blocking solution) was from Biolegend; cell fixable viability dye eFluor 506 was from eBiosciences.

### Cell isolation from various organs

Cells from various organs were isolated as we recently described [40]. Briefly: blood cells were collected through cardiac puncture, and red blood cells were lysed before analysis; lungs were minced and digested in 0.2 mg/ml Liberase TL (Sigma, St. Luis, MO) in 37ºC for 15-30 mins, then filtered and red blood cells were lysed before analysis; intestines were flushed, opened longitudinally, and inner contents removed with the blunt end of scissors, then cut into 0.5-cm fragments, followed by digestion in 100 U/ml collagenase VIII (Sigma) in 37ºC for 1 hour, filtered, and lymphocytes isolated using gradient separation by 40% and 80% Percoll (GE Healthcare, Wilkes-Barre, PA) solutions; perigonadal adipose tissues were minced and digested in 500 U/ml collagenase I (Worthington Biochemical Corp., Lakewood, NJ) in 37ºC for 30 mins, filtered and red blood cells were lysed before analysis. 50-150 U/ml DNase I (Sigma) were added during digestion to reduce cell death triggered by free DNA.

### *In vivo* induction of IL-10-producing T cells by TCR activation

Foxp3^RFP^ IL-10^GFP^ dual reporter mice were injected with 15 µg/mouse anti-CD3ε (145-2C11)intraperitoneally on day 0 and 2, and analyzed on day 4, as previously described [23].

### *Nippostrongylus brasiliensis* (*Nb*) infection

Mice were given 500 L3 *Nb* larvae per mouse through subcutaneous injection, as we previously described [40]. Cells from the lungs were analyzed 7 days post infection (7 dpi).

### House dust mite (HDM)-induced allergic disease model

Mice were given daily intranasal exposures of 10 µg house dust mite (*Dermatophagoides pteronyssinus*) protein extract (XPB82D3A2.5 from Greer) in PBS, for ten consecutive days. Cells from the lungs were analyzed 24 hours post the last treatment.

### Farmer’s lung disease (hypersensitivity pneumonitis) model

Mice were intranasally exposed to 150 µg *Saccharopolyspora rectivirgula* (*SR*, ATCC 29034) extract on 3 consecutive days each week as previously described [41], for four weeks. Cells from the lungs were analyzed on the last day of fourth week.

### Influenza A/WSN/1933 (WSN) infection

Mice were intranasally infected with 1 LD50 (10^4^ PFU) WSN per mouse, as we previously described [40]. Cells from the lungs were analyzed 7 days post infection (7 dpi).

### Differentiation of IL-10-producing T cells

#### Mouse

TCR^+^ Foxp3^RFP−^ CD44^−^ CD62L^+^ splenic naïve T cells were sorted on BD FACS Aria II or Fusion systems (BD Biosciences, San Jose, CA), then cultured with Mitomycin-C (Sigma, 50 µg/ml) treated antigen-presenting cells (APC; *Rag^−/−^* splenocytes) at 1:2 ratio in the presence of anti-CD3ɛ (1 µg/mL), anti-CD28 (1 µg/mL), recombinant murine (rm) IL-27 (R&D Systems, 20 - 25 ng/ml), anti-IFN-γ and anti-IL-12 (10 µg/mL) for 3 days.

#### Human

Human peripheral blood monocytes (PBMCs) were isolated from blood (New York Blood Center, Long Island, NY) using gradient separation in Ficoll-Paque PLUS (GE Healthcare). PBMC were cultured in full RPMI-1640 medium for 30 mins in 37ºC, then non-adherent cells were used to enrich CD4^+^ T cells using anti-human CD4 microbeads (Miltenyl Biotec, San Diego, CA) or CD8^+^ T cells using a human CD8 isolation kit (BioLegend). Adherent cells were treated with Mitomycin-C (Sigma, 50 µg/ml) in 37ºC for 30 mins and used as APCs. Anti-human CD3ɛ (1 µg/ml) and CD28 (CD28.2, eBioscience, 1-3 µg/ml), recombinant human (rh) IL-2 (PeproTech, 200 U/ml), IL-10 (PeproTech, 100 U/ml), IL-27 (R&D System, 25 ng/ml), and IFN-α2b (R&D System, 10 ng/ml) were added to differentiate human IL-10-producing T cells. 3 days after cultures were set up, cells were stimulated with PMA (100 ng/ml, Sigma-Aldrich), Ionomycin (0.5 μM, Sigma), Brefeldin A (5 μg/ml) and GolgiPlug (0.5 μl/ml, BD Biosciences) for 4 hours as we previously described [42], and subjected to surface staining and intracellular staining (see details below).

### Flow cytometry

Surface staining of live cells were done in the presence of Fc block and fixable viability dye. To detect human CD4 in activated T cells, anti-CD4 antibody was added into the intracellular staining panel. For intracellular cytokine staining, cells were fixed with 2% paraformaldehyde (Electron Microscopy Sciences, Hatfield, PA), permeabilized and stained with anti-cytokine antibodies in PBS/0.2% saponin (Sigma). Staining for human transcription factor FOXP3 was performed with a Foxp3 staining buffer kit (eBioscience). Flow cytometry data were acquired on LSRII, FACS Aria II or Fusion systems (BD Biosciences), and analyzed in FlowJo (Tree Star, Ashland, OR). All analyses were performed on fixable viability dye negative singlet population.

### Statistical analysis

Non-parametric Mann-Whitney tests and one-way ANOVA were performed using GraphPad Prism v5.00 (GraphPad, San Diego, CA), with *p* ≤ 0.05 considered statistically significant. “NS” refers to “No Significance”.

## RESULTS

### Co-expression of LAG3 and CD49b marks both IL-10-producing CD4^+^ and CD8+ T cells

LAG3 and CD49b co-expression was previously reported to be a cell surface signature for both mouse and human IL-10-producing CD4^+^ T cells that lack the expression of Foxp3 (also known as type 1 regulatory T cells, Tr1 cells) [15]. We and others have previously reported that co-culturing murine naïve CD4^+^ T cells with antigen presenting cells (APCs) in the presence of anti-CD3, anti-CD28, anti-IFN-γ, anti-IL-12, and IL-27 can efficiently induce the differentiation of Tr1 cells [40; 43; 44], which express high levels of LAG3 and CD49b. Our recent data also demonstrated that this protocol can induce IL-10 production in bulk T cell populations that include both CD4^+^ and CD8^+^ T cells (Fig. 1A). Surprisingly, the resultant IL-10-producing CD8^+^ T cells induced *in vitro* through this protocol also exhibited high levels of LAG3/CD49b co-expression (Fig. 1A, last plot). In fact, IL-10-producing CD8^+^ T cells can express higher levels of LAG3 and CD49b than their IL-10-producing CD4^+^ counterparts induced in the same cell culture (Fig. 1B). These data suggest that co-expression of LAG3 and CD49b is not an exclusive cell surface signature of the Tr1 cells, and may be a shared feature of both IL-10-producing CD4^+^ and CD8^+^ T cells.

**Figure 1:**
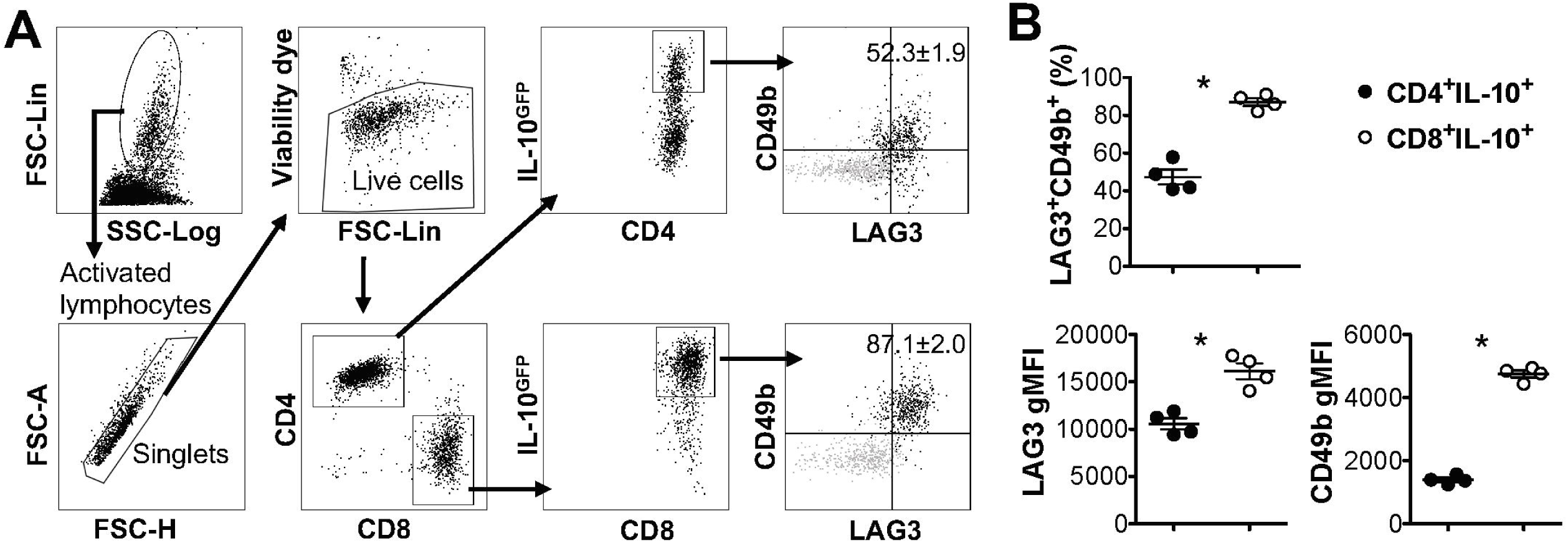
IL-10-producing LAG3^+^ CD49b^+^ T cells include both CD4^+^ and CD8^+^ subsets. All experiments were performed with cells carrying the IL-10^GFP^ reporter system for live cell analysis. Naïve T cells were cultured under IL-10-inducing conditions for 3 days. (**A**) Gating strategy to identify IL-10-expressing CD4^+^ and CD8^+^ T cells, and representative FACS plots showing the co-expression of LAG3/CD49b by IL-10^+^ CD4^+^ or IL-10^+^ CD8^+^ T cells. Grey backgrounds show naïve T cells as negative control for LAG3/CD49b quadrant gating. (**B**) Summary of percentage of LAG3/CD49b double positive population and geometric mean fluorescence intensity (gMFI) of LAG3 and CD49b in IL-10-producing CD4^+^ and CD8^+^ T cells. N = 4. Data represent results of more than three experiments. * *p* ≤ 0.05, by non-parametric Mann-Whitney test. Data presented as Mean ± S.E.M..

### Co-expression of LAG3 and CD49b marks both IL-10-producing Tr1 and Treg cells

IL-10 production can be significantly elevated in the pulmonary mucosa during the late stages of parasitic infection by *Nippostrongylus brasiliensis* (*Nb*), predominantly by CD4^+^ T cells that are LAG3/CD49b double positive [15; 45]. We recently observed that both Foxp3^−^ and Foxp3^+^ T cells are capable of producing IL-10 in the pulmonary tissues post *Nb* infection [40]. To determine whether co-expression of LAG3 and CD49b is exclusive to Foxp3^−^ Tr1 cell subset, we infected IL-10^GFP^/Foxp3^RFP^ dual reporter mice with *Nb*, and analyzed the IL-10-producing T cells. We found that, as previously described, a large majority of the IL-10-producing T cells in the lungs of *Nb*-infected mice express high levels of both LAG3 and CD49b (Fig. 2A&B), and are predominantly CD4^+^ T cells (Fig. 2C). However interestingly, these IL-10-producing LAG3^+^ CD49b^+^ CD4^+^ T cells included both Foxp3^+^ and Foxp3^−^ CD4^+^ T cells subsets (Fig. 2C, last plot; and Fig. 2D). To further determine whether LAG3/CD49b co-expression is correlated with IL-10 and/or Foxp3 expression in CD4^+^ T cells, we compared the percentage of LAG3/CD49b double positive population in Foxp3^−^ IL-10^−^, Foxp3^−^ IL-10^+^, Foxp3^+^ IL-10^−^ and Foxp3^+^ IL-10^+^ CD4^+^ T cells isolated from the lungs of *Nb*-infected mice. We found that regardless of expression of Foxp3, *Nb* infection did not lead to significant up-regulation of LAG3/CD49b co-expression on IL-10^−^ CD4^+^ T cells (Fig. 2E&F). However, the percentage of the LAG3^+^ CD49b^+^ population of IL-10^+^ CD4^+^ T cells is significantly higher than their IL-10^−^ counterparts (Fig. 2F). Moreover, the levels of LAG3 and CD49b expression are similar between the Foxp3^+^ and Foxp3^−^ counterparts of IL-10^+^ CD4^+^ T cells (Fig. 2G). Therefore, IL-10-producing CD4^+^ T cells, regardless of Foxp3 expression, have a high capacity of co-expressing LAG3 and CD49b. Together with the data shown in Fig. 1, these data suggest that co-expression of LAG3 and CD49b is a generic feature of IL-10-producing T cells, including Foxp3^−^ Tr1 cells, Foxp3^+^ Treg cells and CD8^+^ T cells.

**Figure 2:**
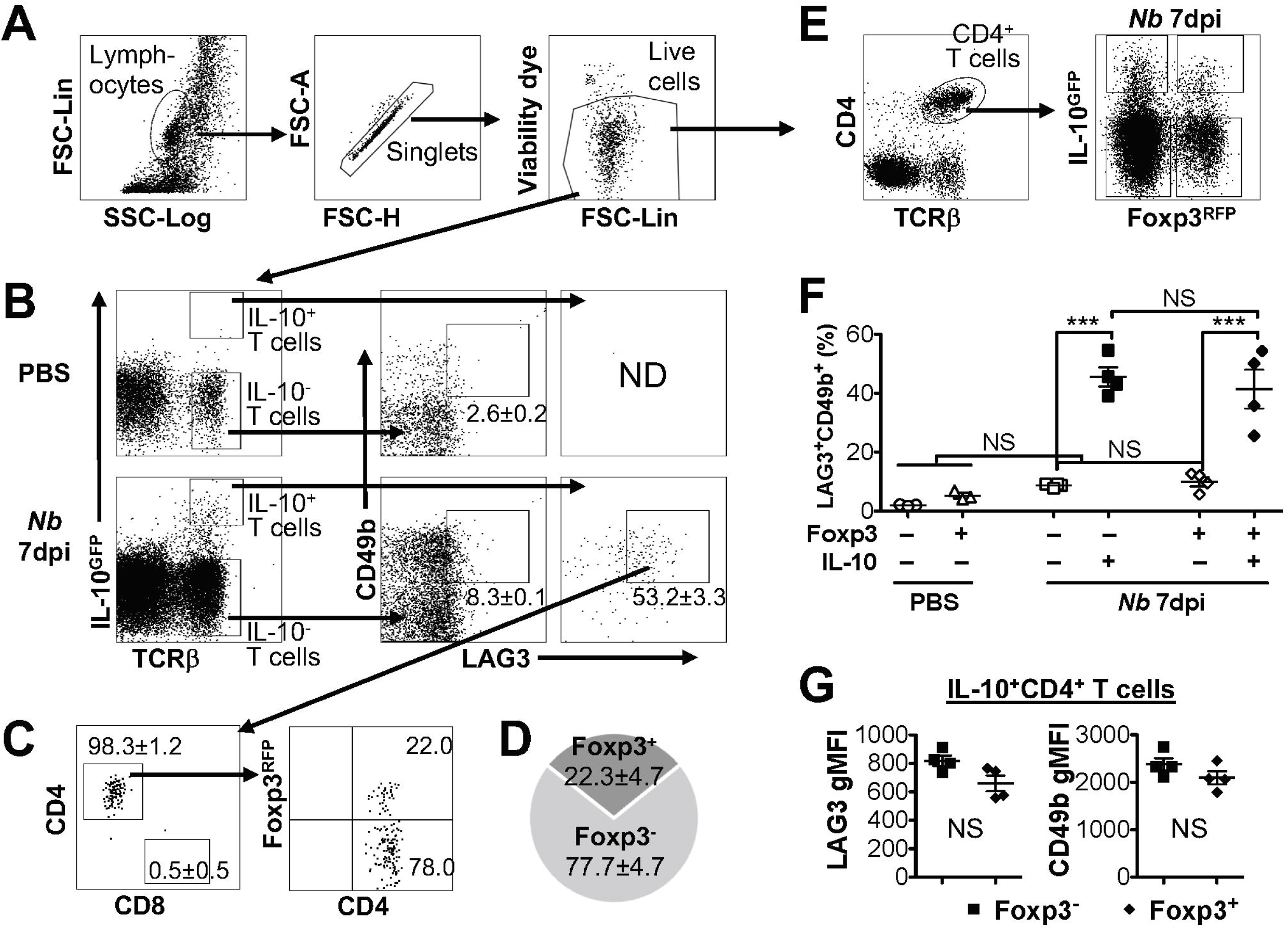
IL-10-producing LAG3^+^ CD49b^+^ CD4^+^ T cells include both Foxp3^+^ and Foxp3-subsets. Mice carrying the IL-10^GFP^/Foxp3^RFP^ dual reporter system were infected with 500 L3 *Nippostrongylus brasiliensis* (*Nb*) (or PBS as control) and lungs analyzed 7 days post infection (dpi). (**A**) Gating strategy to identify live singlet lymphocytes from cells isolated from the lungs of mice for analyses. (**B**) Gating strategy to identify IL-10-producing T cells (with IL-10^−^ T cells as control), and representative FACS plots for LAG3/CD49b co-expression in IL-10^−^ versus IL-10^+^ T cells. (**C**) Representative FACS plots showing that the majority of IL-10^+^ LAG3/CD49b co-expressing T cells in *Nb*-infected mouse lungs are CD4^+^ T cells, including both Foxp3^+^ Treg and Foxp3^−^ Tr1 cells. (**D**) Pie chart summarizing the proportion of Foxp3^+^ versus Foxp3^−^ IL-10-producing LAG3/CD49b double positive CD4^+^ T cells. (**E**) Gating strategy identifying CD4^+^ T cells that are Foxp3^−^ IL-10^−^, Foxp3^−^ IL-10^+^, Foxp3^+^ IL-10^−^ and Foxp3^+^ IL-10^+^. (**F**) Summary of percentage of LAG3/CD49b double positive population of Foxp3^−^ IL-10^−^, Foxp3^−^ IL-10^+^, Foxp3^+^ IL-10^−^ and Foxp3^+^ IL-10^+^ CD4^+^ T cells. (**G**) Summary of gMFI of LAG3 and CD49b in Foxp3^+^ versus Foxp3^−^ IL-10-producing CD4^+^ T cells. N = 4. Data represent results of three experiments. NS = No significance, by non-parametric Mann-Whitney test. Data presented as Mean ± S.E.M..

### The composition of LAG3^+^ CD49b+ IL-10-producing T cells differs in different disease models

IL-10 plays an essential role in pulmonary inflammatory diseases, reported in multiple murine models of lung diseases, including allergic asthma [46], hypersensitivity pneumonitis (HP) [47] and influenza pneumonia [20]. We examined whether the expression of LAG3 and CD49b would differ based on the inflammatory response in three mouse models of lung inflammation. Mice carrying the IL-10^GFP^/Foxp3^RFP^ dual reporters were exposed intranasally to house dust mite (HDM) protein extract (as a model of allergic asthma), *Saccharopolyspora rectivirgula (SR)* (as a model of HP/farmers’ lung disease), or infected intranasally with WSN/flu virus (as a model of influenza infection). We observed significant percentages of IL-10-producing T cells in the lung tissue of mice exposed HDM (Fig. 3A), *SR* (Fig. 3C), or WSN/flu virus (Fig. 3E). These IL-10-producing T cells all co-expressed high levels of LAG3 and CD49b, and include Foxp3^+^ CD4^+^, Foxp3^−^ CD4^−^ and CD8^+^ subsets in all disease models analyzed (Fig. 3). However, the relative proportions of IL-10-producing Foxp3^+^ CD4^+^, Foxp3^−^ CD4^−^ and CD8^+^ subsets differed in the various disease models (Fig. 3). In contrast to composition of IL-10-producing LAG3^+^ CD49b^+^ T cells induced by *Nb* infection, in which Foxp3^−^ CD4^+^ subset is the majority (72% in Fig. 2B), in HDM-induced allergic asthma model, the largest subset of the IL-10-producing LAG3^+^ CD49b^+^ T cells in lungs are Foxp3^+^ CD4^+^ T cells (60%), followed by Foxp3^−^ CD4^+^ T cells (32%), while CD8^+^ T cells are only around 1.4% (Fig. 3B). In the *SR*-triggered farmer’s lung disease model, Foxp3^+^ CD4^+^ T cells are the largest majority of the IL-10-producing LAG3^+^ CD49b^+^ T cell subset in the lungs, however, there are similar percentages of Foxp3^−^ CD4^+^ and CD8^+^ T cells (16% each) (Fig.3D). Strikingly in the murine model of influenza infection, CD8^+^ T cells are the largest subset of the IL-10-producing LAG3^+^ CD49b^+^ T cells in the lungs (86%), followed by Foxp3^−^ CD4^+^ T cells, while Foxp3^+^ CD4^+^ are a minority (Fig. 3F). These data compared the composition of IL-10-producing LAG3^+^ CD49b^+^ T cells in various murine models of pulmonary inflammatory diseases. Along with the model of parasitic infection shown in Fig. 2, our data suggest that co-expression of LAG3 and CD49b marks all IL-10-producing T cells in the pulmonary system, and relative abundance of the marked T cell subsets is dependent on the type of immune response as shown in the disease models.

**Figure 3:**
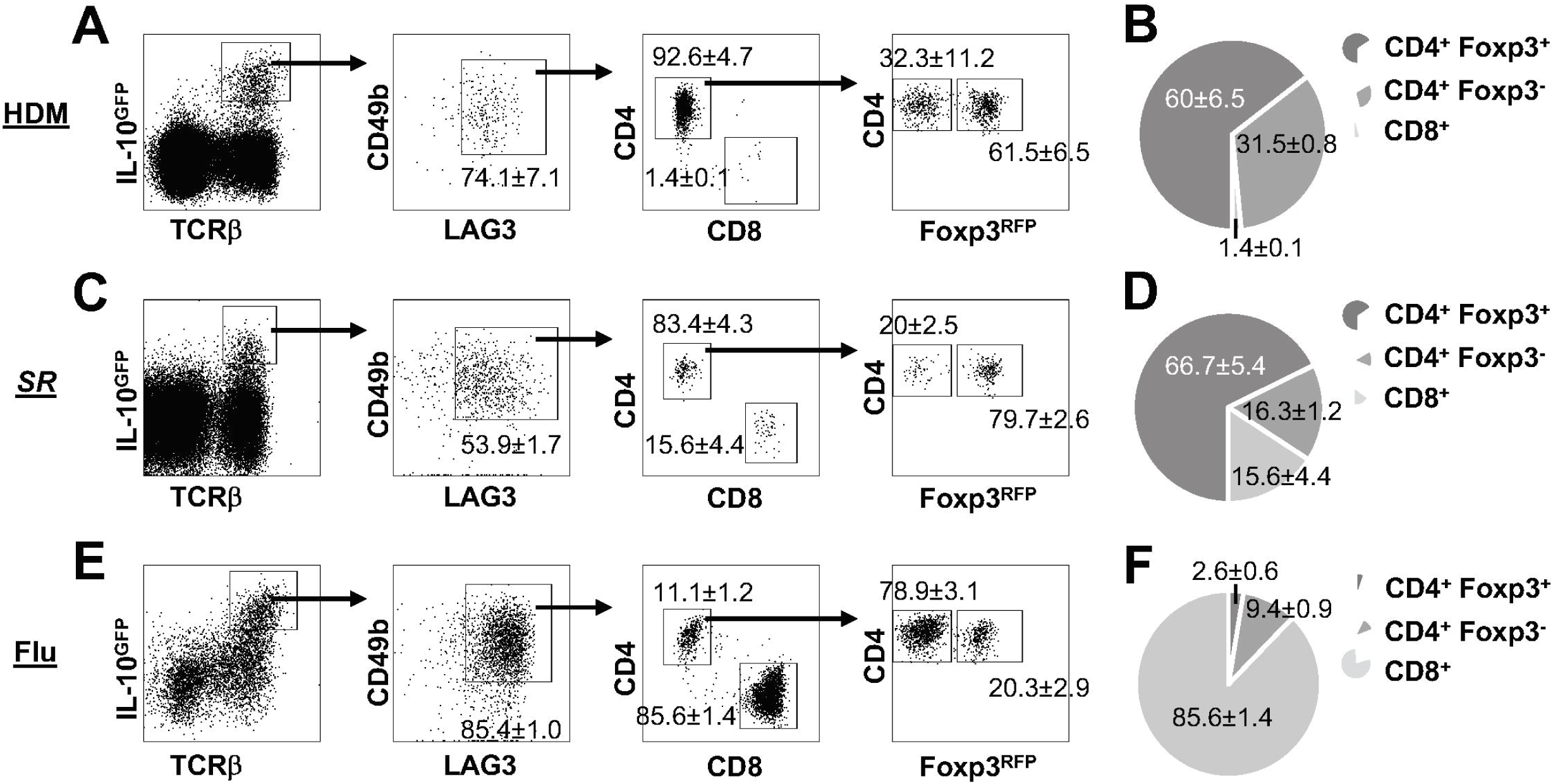
Composition of IL-10-producing LAG3^+^ CD49b^+^ T cells in lung inflammatory and infectious disease models. All experiments were performed using mice carrying the IL-10^GFP^/Foxp3^RFP^ dual reporter system, and cells were isolated from the lungs and gated on live singlets for analysis as shown in Fig. 2A. Representative FACS plots of LAG3/CD49b co-expression by IL-10-producing cells, and among these cells, ratio of CD4^+^ versus CD8^+^ T cells, and Foxp3^+^ versus Foxp3^+^ CD4^+^ subpopulations are shown, followed by pie chart showing the proportions of CD4^+^ Foxp3^+^, CD4^+^ Foxp3^−^, and CD8^+^ T cells in IL-10-producing LAG/CD49b double positive T cell population. Mice were exposed intranasally to: (**A-B**) House dust mite (HDM) protein extract for 10 consecutive days, and analyzed 24 hours post the last treatment; (**C-D**) *Saccharopolyspora rectivirgula (SR)* for 3 consecutive days every week for 4 weeks, and analyzed in the end of the fourth week; (**E-F**) influenza A (WSN) viruses, and analyzed 7 dpi. N ≥ 3, combined from three independent experiments. Data presented as Mean ± S.E.M..

### The composition of LAG3^+^ CD49b+ IL-10-producing T cells differs in different organs

As discussed above, we demonstrated that co-expression of LAG3 and CD49b is a generic feature of IL-10-producing T cells *in vivo* in the pulmonary tissues under multiple inflammatory conditions (Figs. 2–3). To determine whether this feature is applicable to IL-10-producing cells in other organs, we injected IL-10^GFP^/Foxp3^RFP^ dual reporter mice with an anti-CD3ε antibody that has been shown to stimulate pronounced IL-10 production by T cells through TCR activation *in vivo* [23; 40]. We analyzed IL-10-producing T cells in blood, lymph nodes (LN), lung, fat and small intestine, found that co-expression of LAG3 and CD49b marked a portion of the IL-10-producing T cells following TCR activation *in vivo*, which again included Foxp3^+^ CD4^+^, Foxp3^−^ CD4^+^ and CD8^+^ T cell subsets (Fig. 4A) in all organs analyzed. An interesting note is that the relative abundance of Foxp3^+^ CD4^+^, Foxp3^−^ CD4^+^ and CD8^+^ subsets among the IL-10-producing LAG3^+^ CD49b^+^ T cells vary significantly in different organs of the same mice (Fig. 4B). Upon TCR activation *in vivo*, Foxp3^+^ CD4^+^ T cells are the major population that are IL-10^+^ LAG3^+^ CD49b^+^ in the blood, lymph nodes and lungs, while CD8^+^ T cells are the majority of IL-10^+^ LAG3^+^ CD49b^+^ T cells in the perigonadal fat and small intestine (Fig. 4B). These data suggest that co-expression of LAG3/CD49b marks all three IL-10-producing T cell subsets in multiple organs and the relative abundance of the Foxp3^+^ CD4^+^, Foxp3^−^ CD4^+^ and CD8^+^ T cell subsets in IL-10-producing LAG3^+^ CD49b^+^ T cells is dependent on the anatomical location of the cells.

**Figure 4:**
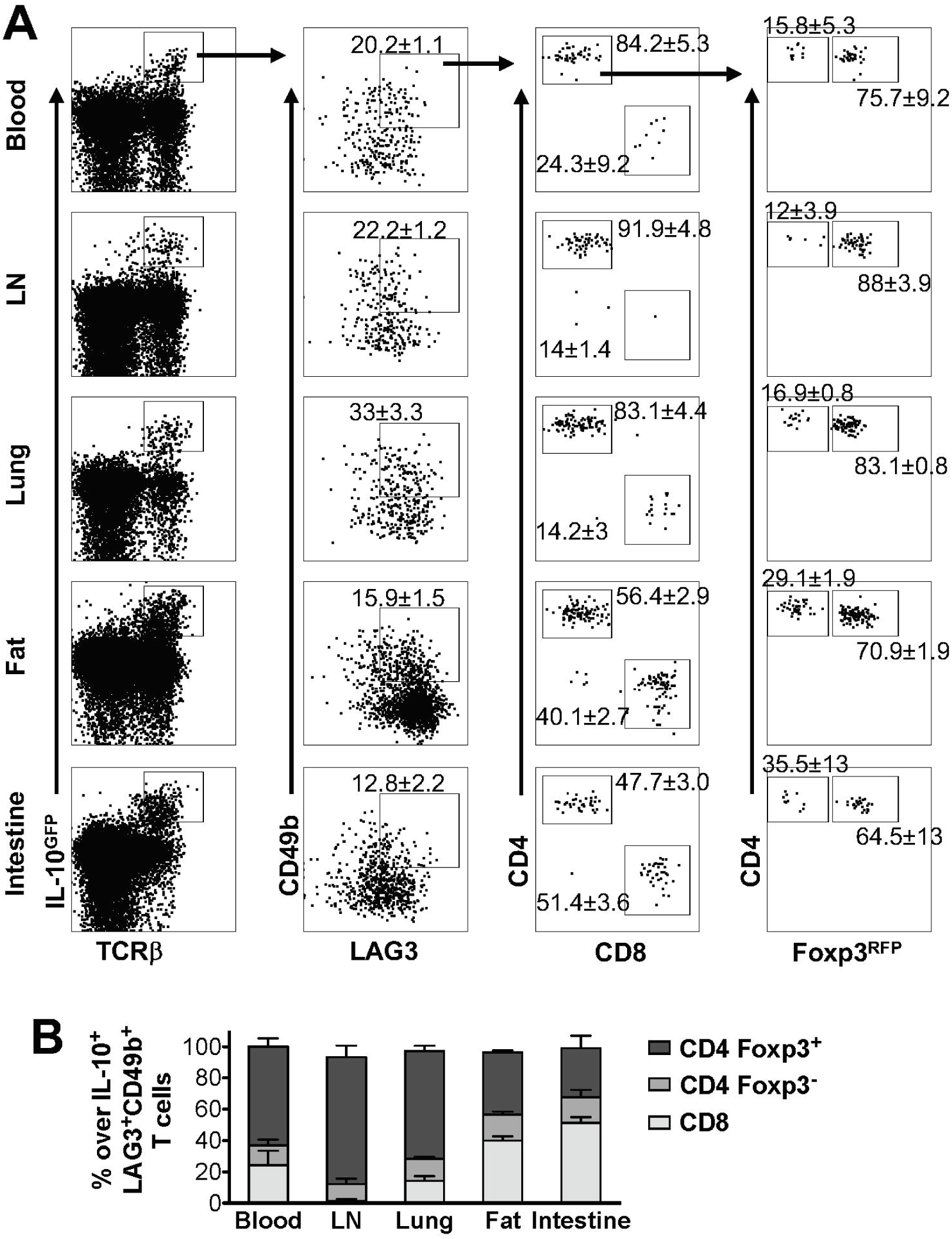
Composition of IL-10-producing LAG3^+^ CD49b^+^ T cells in various organs *in vivo* following TCR activation. Mice carrying the IL-10^GFP^/Foxp3^RFP^ dual reporter system were injected with CD3 antibody on day 0 and 2, and analyzed on day 4. Live singlet cells from the indicated organs were gated for analysis. (**A**) Representative FACS plots showing LAG3/CD49b co-expression by IL-10-producing cells, and among these cells, proportion of CD4^+^ versus CD8^+^ T cells, and Foxp3^+^ versus Foxp3^+^ CD4^+^ subpopulations. (**B**) Stacked bar chart summarizing the percentages of Foxp3^+^ CD4^+^, Foxp3^−^ CD4^+^, and CD8^+^ T cells in IL-10-producing LAG/CD49b double positive T cell population in the indicated organs. N = 3. Data represent results of three experiments, and are presented as Mean ± S.E.M..

### Human IL-10-producing CD4^+^ and CD8+ T cells exhibit LAG3+CD49b+ phenotype

In murine models, we demonstrated that co-expression of LAG3 and CD49b marks IL-10-producing Foxp3^+^ CD4^+^, Foxp3^−^ CD4^+^ and CD8^+^ T cells under different inflammatory conditions in the lungs (Fig. 2–3), as well as in different organs when TCR is activated *in vivo* (Fig. 4). To further determine whether different subsets of IL-10-producing T cells exhibit the shared feature of co-expression of LAG3 and CD49b in human, we isolated CD4^+^ and CD8^+^ T cells from human peripheral blood and cultured them under IL-10-inducing conditions. We found that as in the mouse, human IL-10-producing FOXP3^+^ CD4^+^, FOXP3^+^ CD4^+^ and CD8^+^ subsets all up-regulated both LAG3 and CD49b expression, with a significant LAG3/CD49b double positive population (Fig. 5).

**Figure 5:**
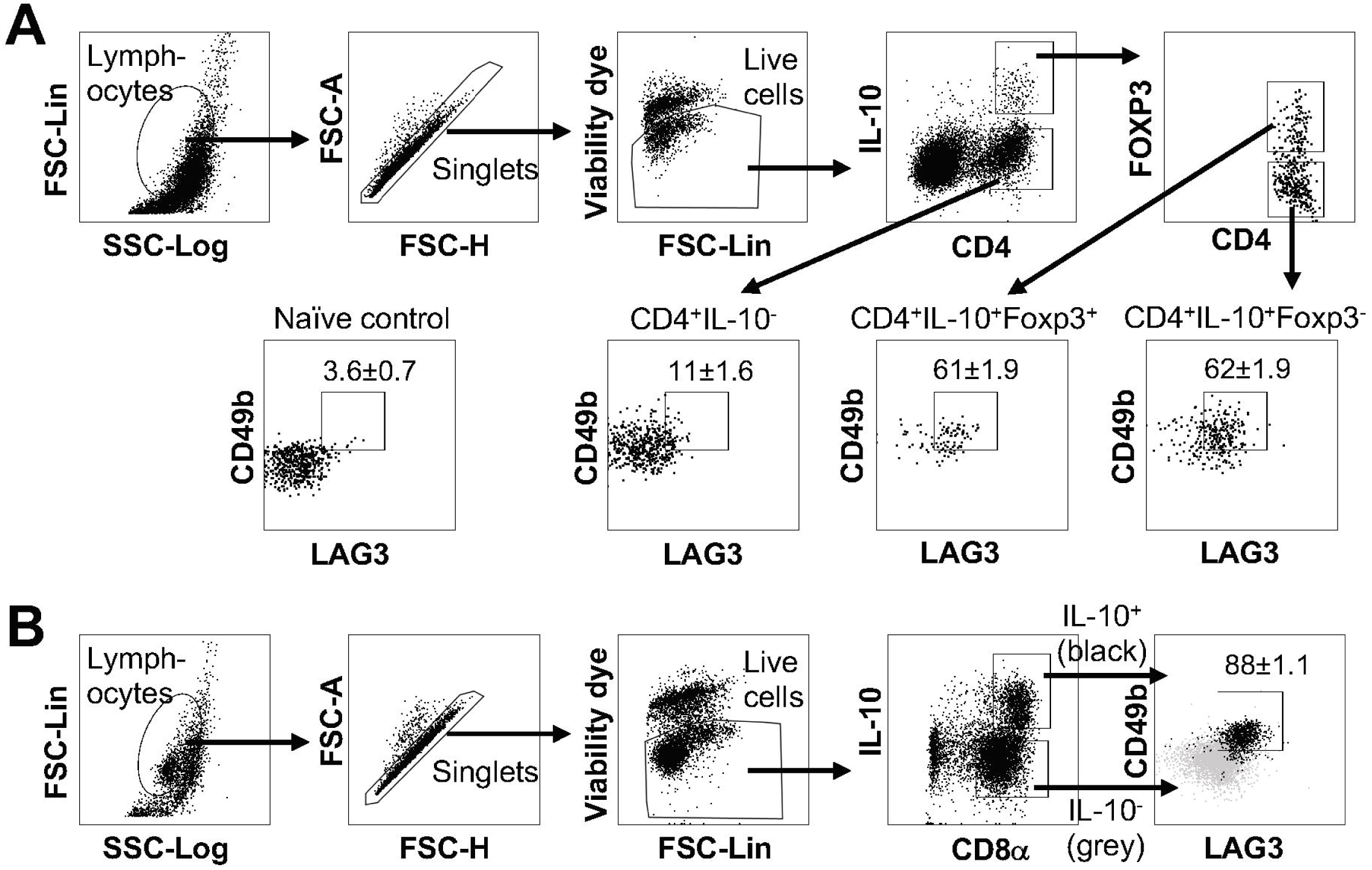
Human IL-10-producing Treg, Tr1 and CD8^+^ T cells exhibit LAG3^+^ CD49b^+^ phenotype. CD4^+^ or CD8^+^ T cells isolated from human peripheral blood mononuclear cells were cultured under IL-10-inducing conditions; cells were stimulated and subjected to intracellular staining. (**A**) Representative FACS plots showing gating strategy to identify IL-10^−^ CD4^+^ T cells and IL-10-producing FOXP3^+^ and FOXP3^−^ CD4^+^ T cells and percentage of LAG3/CD49b double positive subset in these cells. Naïve CD4^+^ T cells were use as negative control to set the LAG3/CD49b double positive gate. N = 4. Data represent results of more than three independent experiments. (**B**) Representative plots showing gating strategy and LAG3/CD49b co-expression for IL-10-producing CD8^+^ T cells. Grey backgrounds show IL-10^−^ CD8^+^ T cells as control. N = 6. Data represent results of two independent experiments. Data presented as Mean ± S.E.M..

## DISCUSSION

Our data presented in this report demonstrated that, in contrast to previously reported [15], co-expression of LAG3 and CD49b is not an exclusive cell surface signature of Foxp3^−^ IL-10^high^ Tr1 cells in human and mouse. In addition, we find that the IL-10-producing LAG3^+^ CD49b^+^ T cell population is composed of Foxp3^+^ CD4^+^, Foxp3^−^ CD4^+^, and CD8^+^ T cell subsets, and this composition varies depending on disease conditions and anatomical locations of the cells. The significance of this work is emphasized due to the importance of identifying Foxp3^−^ Tr1 cells, including under clinical conditions, and our findings urge caution in the use of LAG3 and CD49b to uniquely identify these cells.

Using Tr1 cell clones derived from human naïve CD4^+^ T cells and purified by an IL-10 secretion assay, Gagliani, Roncarolo and colleagues identified co-expression of LAG3, CD49b and CD226 as cell surface signatures of IL-10-producing CD4^+^ T cells, and demonstrated that co-expression of LAG3 and CD49b is sufficient to distinguish Foxp3^−^ IL-10^high^ Tr1 cells from T helper and/or regulatory subsets that expressed lower levels of or no IL-10 in both mouse and human [15]. However, it was recently reported that IL-10-producing T cells derived from human CD4^+^ memory T cells exhibit low levels of surface expression of LAG3 and CD49b [48], suggesting that the pattern of co-expression of LAG3 and CD49b may vary in human IL-10-producing CD4^+^ T cells, which may be associated with whether they were derived from naïve precursors versus memory cells. Indeed, co-expression of LAG3 and CD49b may not be able to mark all IL-10-producing Tr1-like cells, but may serve to help eliminate those T cells lineages that are not capable of producing IL-10 for potential clinical interest. Whether co-expression of LAG3 and CD49b would allow efficient recovery of IL-10^high^ T cells may dependent on the proportion of IL-10^high^ T cells that are co-expressing LAG3 and CD49b, which may differ dependent on whether the relevant IL-10-producing cells had different origins and/or underwent different activation regimes.

Both LAG3 and CD49b can individually be up-regulated in activated T cells, regardless of their production of the anti-inflammatory IL-10 or pro-inflammatory cytokines [31; 32; 33; 35; 36; 37]. Co-expression of LAG3 and CD49b is more restricted to IL-10-producing subsets, as previously described in Foxp3^−^ CD4^+^ T cells [15]. And our data here demonstrated that this is a generic feature of IL-10-producing cells, including Foxp3^−^ CD4^+^, Foxp3^+^ CD4^+^ and CD8^+^ subsets. The co-occurrence of IL-10 production and LAG3^+^ CD49b^+^ may be explained in two ways. The first explanation is that IL-10 stimulation through autocrine and/or paracrine may induce the expression of LAG3 and CD49b in the stimulated cells, therefore, IL-10-capturing T cells (cells detected by Roncarolo’s group in ref [15]) exhibited high levels of LAG3/CD49b co-expression. This hypothesis would place IL-10 up-stream of LAG3 and CD49b expression, which is unlikely, as recent studies by Flavell and Huber’s groups showed that IL-10 receptor is dispensable for Tr1 cell differentiation, including the induction of LAG3/CD49b double positive feature; instead, IL-10 receptor signaling is critical for maintaining the cell fate commitment and functional performance of the differentiated Tr1 cells [49]. Therefore, it is more likely that LAG3 and CD49b signaling pathways function cooperatively to activate the expression of IL-10. This hypothesis is more reasonable, given our data that LAG3 and CD49b co-expression is a generic feature of IL-10 producing cells in multiple subsets. LAG3 is a structural homology of CD4 molecule, and can bind to MHC class II in higher affinity than CD4 [29; 30]. In unstimulated T cells, LAG3 is retained in the intracellular compartment and degraded in the lysosome; upon T cell activation, LAG3 traffics from the lysosomal compartment to the cell surface through a protein kinase C (PKC) dependent pathway [50]. CD49b is the integrin α2 subunit and plays a critical role in cell-cell interaction and cell-surface adhesion [51]. The CD49b pathway activates multiple downstream effector signaling pathways, among which there is the RAS/MAPK signaling cascade [52; 53]. We recently reported that the RAS/MAPK signaling pathway functioning downstream of the TCR is indispensable for IL-10 production by Foxp3^−^ CD4^+^ cells, through activation of the expression of the transcription factor interferon regulatory factor 4 (IRF4) [40]. PKC can regulate RAS signaling to downstream effectors, and can activate MAPK signaling in both RAS-dependent and independent manners [54; 55]. The interplay between PKC and RAS may regulate signals that connect LAG3 and CD49b downstream pathways, leading to up-regulation of IL-10. Given the complexity of these pathways, comprehensive understanding of the mechanism(s) underlying the co-expression of these markers with IL-10 will require significantly more in-depth analyses. Regardless of the mechanism, our findings indicate that LAG3/CD49b co-expression does not uniquely identify Foxp3^−^ Tr1 cells, but is a more general indicator of IL-10 production in T cell lineages.

Despite the shared feature of co-expression of LAG3 and CD49b by IL-10-producing Foxp3^+^ CD4^+^, Foxp3^−^ CD4^+^ and CD8^+^ T cells, we also observed interesting discrepancies in the proportional composition of these three IL-10^high^ T cell subsets that are all LAG3/CD49b double positive in the lung mucosa of different pulmonary inflammatory disease models, as well as in different anatomical locations in the same mice upon TCR activation *in vivo*. For example, parasite infection (*Nb*) and mite allergen (HDM) induced predominantly IL-10-producing LAG3^+^ CD49b^+^ T cells that are CD4^+^ with very few that are CD8^+^. In *Nb* the infection model, the majority of IL-10^+^ LAG3^+^ CD49b^+^ CD4^+^ T cells are Foxp3^−^ Tr1 cells, while in HDM-treated mice, Foxp3^+^ Treg cells are the majority. In the bacterial exposure (*SR*) and viral infection (Flu) models, we observed an increased proportion of IL-10^+^ LAG3^+^ CD49b^+^ T cells that are CD8^+^, which is the predominant population in the Flu model (Fig. 3F). This discrepancy in the composition of IL-10-producing LAG3^+^ CD49b^+^ T cells may be due to the difference of the microenvironment in which the IL-10-producing cells are being induced. Factors that may affect the different composition of the IL-10^high^ LAG3^+^ CD49b^+^ T cells may include the type of immune response, abundance and affinity of the TCR ligands, the cytokines induced and orchestrated by the stimuli, and the nutritional microenvironments that favor different subsets of the T cells. For example, type I IFN can facilitate the preferential induction of IL-10-producing effector CD8^+^ T cells thorough inducing and sustaining expression of the IRF4 and Blimp1 transcription factors [56]. Type I interferon is significantly elevated during influenza infection [57] but much less so by HDM exposure [58].

Another possible orchestrator could be the cytokine IL-27, which has been reported to be directly required for IL-10 induction in CD8^+^ T cells [59] and CD4^+^ Foxp3^−^ [60] but not CD4^+^ Foxp3^+^ T cell subsets [61]. Transcription factors and their interacting molecular networks that exhibit differential functions for IL-10 induction in different T cell lineages might provide an answer as well. For example, Blimp-1 is indispensable for IL-10 induction in Foxp3^+^ and Foxp3^−^ CD4^+^ T cells [62; 63], as well as in CD8^+^ T cells [64]. AhR interaction with cMAF is critical in IL-10 induction in Foxp3^−^ Tr1 cells in response to IL-27-supplemented environment, but is insufficient in inducing IL-10 production in Foxp3^+^ Treg cells under the same conditions [43]; AhR expression was not critical in IL-10-producing CD8^+^ T cells either [65], suggesting that AhR signaling has multifaceted function in regulating the level of IL-10 expression in different T cell lineages. A more comprehensive understanding of the T cell-intrinsic molecular features that are shared or distinct among the IL-10-producing CD8^+^, Foxp3^−^ CD4^+^ and Foxp3^+^ CD4^+^ T cells awaits further investigation.

Our data reported here demonstrate that co-expression of LAG3 and CD49b marks IL-10^high^ T cell subsets that are Foxp3^+^ CD4^+^, Foxp3^−^ CD4^+^ or CD8^+^ in both human and mouse, and thus does not uniquely identify Foxp3^−^ Tr1 cells. However, this finding does not negate the feasibility of utilizing co-expression of LAG3 and CD49b in marking a broader range of immunosuppressive IL-10^high^ T cell populations that have potential therapeutic effects for clinical application. Further investigations are required to determine the levels of regulatory and pro-inflammatory cytokine production in the bulk LAG3^+^ CD49b^+^ T cells, and to determine and compare the ability of the Foxp3^+^ CD4^+^, Foxp3^−^ CD4^+^ and CD8^+^ subsets of the LAG3^+^ CD49b^+^ T cells in suppressing effector immunity and inflammation *in vivo*.

## Acknowledgements

We thank A. Redko for animal care and L. Zhang for technical assistance; Dr. E. Tait Wojno for *N. brasiliensis*; and Drs. D. Topham, G. Whittaker, and M. Straus for influenza A virus. This work was supported in part by grants from the National Institutes of Health (AI120701, AI138570 and AI126814 to A.A.; AI129422 and AI138497 to A.A. and W.H., and AI137822 to W.H.), a Careers in Immunology Fellowship from the American Association of Immunologists (to W.H.), a Pilot Grant from the Center for Experimental Infectious Disease Research (funded by NIH P30GM110760) and the Faculty Development Program of the Louisiana State University (to W.H.), and the Talent Program of The Third Affiliation Hospital of Sun Yat-sen University (555 to W.H. and S.G.Z.).

## Author Contributions

W.H. and A.A. conceived research, designed experiments, analyzed and interpreted data, and wrote the manuscript; W.H., S.S., and C.C. performed experiments; S.G.Z. contributed reagents and intellectual input.

## Abbreviations Used

Nb: Nippostrongylus brasiliensis
Tr1 cell: Type 1 regulatory T (CD4^+^ TCR^+^ Foxp3^−^ IL-10^+^) cell
Treg cell: Foxp3-expressing regulatory T cell
HDM: house dust mite
SR: Saccharopolyspora rectivirgula
WSN: Influenza A/WSN/1933 (H1N1)

